# HBEGF^+^ macrophages identified in rheumatoid arthritis promote joint tissue invasiveness and are reshaped differentially by medications

**DOI:** 10.1101/525758

**Authors:** David Kuo, Jennifer Ding, Ian Cohn, Fan Zhang, Kevin Wei, Deepak Rao, Cristina Rozo, Upneet K. Sokhi, Accelerating Medicines Partnership RA/SLE Network, Edward F. DiCarlo, Michael B. Brenner, Vivian P. Bykerk, Susan M. Goodman, Soumya Raychaudhuri, Gunnar Rätsch, Lionel B. Ivashkiv, Laura T. Donlin

## Abstract

Macrophages tailor their function to the signals found in tissue microenvironments, taking on a wide spectrum of phenotypes. In human tissues, a detailed understanding of macrophage phenotypes is limited. Using single-cell RNA-sequencing, we define distinct macrophage subsets in the joints of patients with the autoimmune disease rheumatoid arthritis (RA), which affects ~1% of the population. The subset we refer to as HBEGF^+^ inflammatory macrophages is enriched in RA tissues and shaped by resident fibroblasts and the cytokine TNF. These macrophages promote fibroblast invasiveness in an EGF receptor dependent manner, indicating that inflammatory intercellular crosstalk reshapes both cell types and contributes to fibroblast-mediated joint destruction. In an *ex vivo* tissue assay, the HBEGF^+^ inflammatory macrophage is targeted by several anti-inflammatory RA medications, however, COX inhibition redirects it towards a different inflammatory phenotype that is also expected to perpetuate pathology. These data highlight advances in understanding the pathophysiology and drug mechanisms in chronic inflammatory disorders can be achieved by focusing on macrophage phenotypes in the context of complex interactions in human tissues.

**One Sentence Summary:** A newly identified human macrophage phenotype from patients with the autoimmune condition RA is found to promote joint tissue invasiveness and demonstrates variable sensitivities to anti-inflammatory medications used to treat the disease.

## Introduction

Macrophage plasticity provides tailored homeostatic, immunologic and reparative mechanisms in a wide-range of tissues (*1, 2*). Their transcriptional, epigenetic and functional versatility allow macrophages to conform to tissue- and disease-specific factors, resulting in phenotypes indicative of the type of tissue and physiologic state (*3–9*). While macrophages are a unifying feature in chronic human diseases such as atherosclerosis, autoimmunity and granulomas (*2, 10, 11*), little is known about macrophage phenotypes in the context of human tissue pathology—particularly at the single-cell level. Furthermore, while *in vitro* studies have provided valuable insights into the range of macrophage polarization states (*12, 13*), the relevance of these well-characterized responses has been difficult to document in human tissues.

A precise understanding of human tissue macrophages may enable more effective therapeutic decisions for inflammatory diseases, where it has often been difficult to discern which molecular pathways to target. For example, in inflammatory bowel diseases cytokines such as IL-17 and IFN-γ have been implicated in the pathophysiology, yet blockade of these factors have produced variable results and in some cases worsens symptoms, whereas anti-TNF therapies are commonly effective (*14*). In the autoimmune disease rheumatoid arthritis (RA) nonsteroidal anti-inflammatory drugs (NSAIDS) treat pain and components of inflammation, but for unclear reasons do not curb joint erosion (*15*), whereas anti-TNF therapies have proven highly effective on both fronts. Macrophages, as innate immune cell types common to tissues that are heavily influenced by microenvironmental factors, could serve in affected tissues as indicators of disease pathways and as targets of tailored treatments. Furthermore, in cancer, new therapeutic strategies not only disrupt support of tumor growth and metastasis by macrophages, but aim to repolarize them towards pro-resolution anti-tumor states (*16, 17*). To develop such therapeutics for autoimmune and inflammatory conditions, an in-depth classification is needed for tissue macrophages and how medications redirect them, amidst the complexity of tissue and disease signals, towards beneficial or a differing yet pathologic state.

In the chronically inflamed RA joint tissue, macrophages are understood to be a source of TNF, a well-established driver of RA (*18–21*). However, the precise nature and variety of macrophages within the RA synovium is not yet defined, nor is the cumulative impact of surrounding intercellular interactions and medication-induced perturbations on macrophage responses in RA tissue pathology.

## RESULTS

### Single-cell RNA sequencing detects HBEGF^+^ inflammatory macrophages in RA synovial tissue

To define the spectrum of macrophage phenotypes in a human tissue affected by autoimmunity, as reported (*22*), we sorted synovial cells from ten RA patients and two osteoarthritis (OA) disease control patients by CD14^+^ cell surface protein expression and applied single-cell RNA sequencing (scRNA-seq, CEL-Seq2). After stringent quality control filtering, 940 CD14^+^ single cells clustered into four major CD14^+^ synovial cell subsets based on Canonical Correlation Analysis (CCA) as described (*22*)(Fig. 1A). Cells from all four clusters expressed genes that define the myeloid lineage, such as *CD68*, *CD163* and *C1QA* and based on their high levels of *CD14* (Fig. S1A), we designated these cells macrophages rather than dendritic cells (*7, 23*). Clusters 1 and 2 contained the majority of the CD14^+^ synovial cells (45% and 30%, respectively) and the largest number of genes that distinguished them from the other clusters, such as *PLAUR* and *HBEGF* for Cluster 1 (referred to as the ‘Cluster 1 HBEGF^+^) and *ADORA3* and *MERTK* for Cluster 2 (Fig. 1B, Fig. S1B, 125 and 193 genes >2-fold, respectively). Cluster 3 appeared less well defined by positive markers, whereas Cluster 4 had robust markers including IFN-target genes such as *IFI6* and *IFI44L* and herein is referred to as the ‘IFN/STAT’ cluster (Fig. 1B, Fig. S1B).

**Fig. 1.**
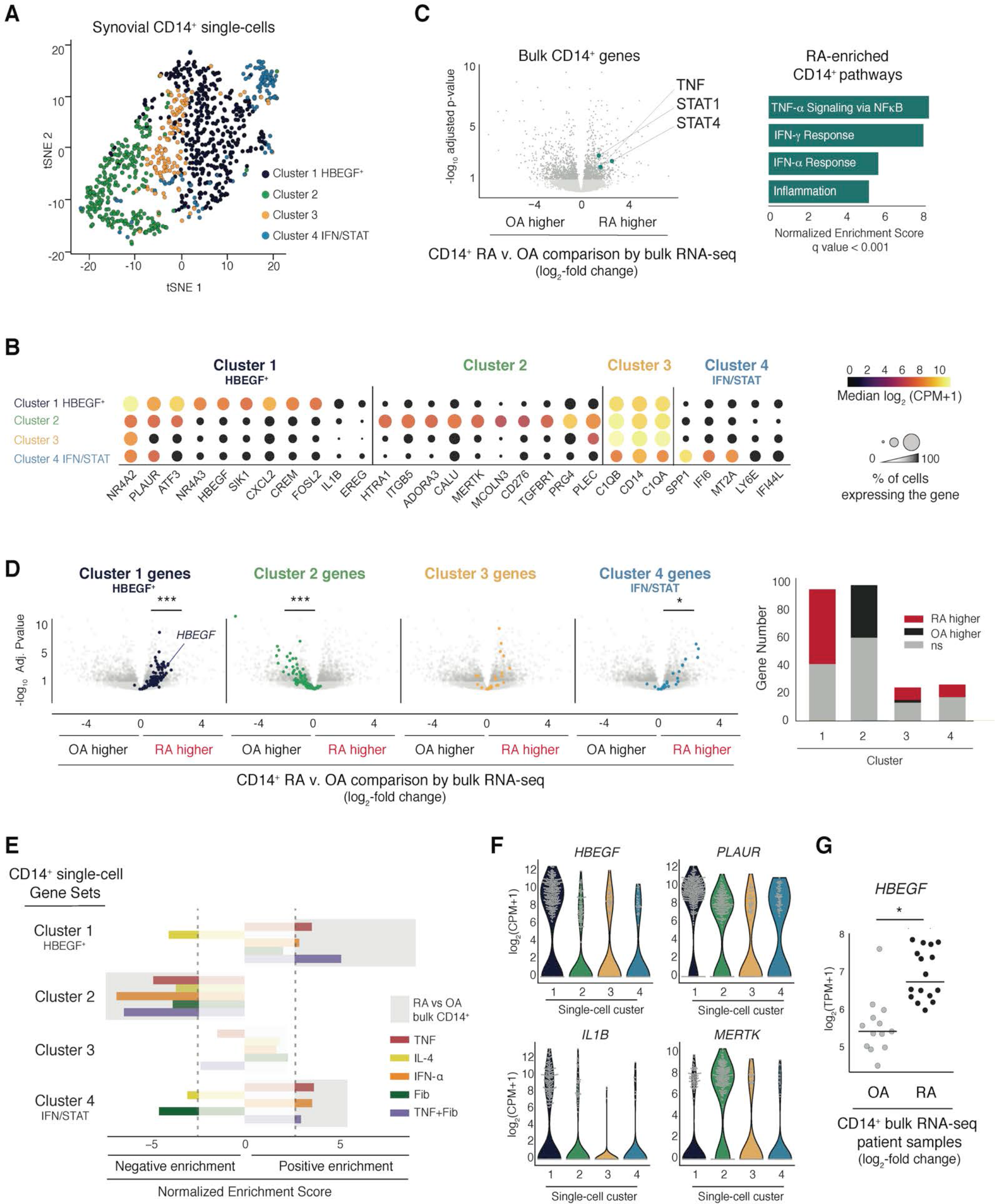
HBEGF^+^ inflammatory macrophage identification in RA joints by single-cell RNA-sequencing. (**A**) Synovial CD14^+^ single-cell RNA-seq clusters (940 cells) identified by Canonical Correlation Analysis (*22*). (**B**) CD14^+^ single-cell cluster marker genes. Median expression indicated by color and percentage of expressing cells indicated by size. (**C**) Differential gene expression for bulk CD14^+^ synovial cells from RA (n=16) versus OA (n=13) patients plotted as *log*_2_ fold-change with *−log*_10_ FDR adjusted p-value; dark grey <0.1. Right: Positively enriched pathways in RA bulk CD14^+^ cells. (**D**) CD14^+^ single-cell cluster genes (up to 100) highlighted on the bulk RA v. OA plot from b (Bonferroni corrected p-value < 0.1, expressed in ≥30% of cells). Hypergeometric test: ***,* represent p <10^−6^, <10^−3^, respectively. Right: Number of cluster genes higher in RA or OA bulk comparison or not significant (ns) (FDR adjusted p-value < 0.1). (**E**) GSEA using the CD14^+^ single-cell markers as Gene Sets and ranked gene lists from human blood-derived macrophages exposed to various stimuli (colored bars) or the bulk CD14^+^ RA v. OA analysis (background grey bars). Normalized enrichment score; |NES| >2.5 were significant at FDR adjusted p <0.001. (**F**) Gene expression level for each cell, plotted as *log*_2_ counts per million (CPM)+1. (**G**) *HBEGF* expression in patient CD14^+^ bulk populations, plotted as *log*_2_ transcripts per million (TPM)+1. n=16 RA, 13 OA samples. *, Bonferroni corrected p value <10^−3^.

To estimate the abundance of the macrophage clusters in RA tissue, we sorted CD14^+^ synovial cell populations from 18 RA and 14 OA patients and analyzed them using bulk RNA-seq (~1,000 CD14^+^ cells from each patient) as reported (*22*). We detected 1,726 genes with distinct expression patterns between the disease states (FDR adjusted p value <0.1). Enriched in RA CD14^+^ cell populations, we detected elevated levels of genes and pathways that associate with RA disease such as *TNF*, *STAT1* and *STAT4* and the response to TNF, Interferon and Inflammation (Gene Set Enrichment Analysis (GSEA)) (Fig. 1C, left and right panel, respectively) (*24–27*). Overlaying the RA versus OA comparison with markers from each of the single-cell clusters, we found that genes defining Cluster 1 HBEGF^+^ and Cluster 4 IFN/STAT were consistently more abundant in RA CD14^+^ populations, suggesting an enrichment of these cell subsets in RA (Fig. 1D). Cluster 2 genes were in higher ratios in OA tissues, while Cluster 3 markers showed no consistent association with either disease (Fig. 1D). Furthermore, RA CD14^+^ populations were positively enriched in genes sets from Cluster 1 HBEGF^+^ and Cluster 4 IFN/STAT while negatively enriched in Cluster 2 genes (GSEA, FDR adjusted p-value <0.001)(Fig. 1E, light grey bars).

To identify factors from the microenvironment that shape synovial macrophage phenotypes, the synovial CD14^+^ single-cell gene sets were compared to human blood-derived macrophages activated by diverse stimuli (Fig. 1E, colored bars, ranked gene lists). Cluster 1 HBEGF^+^ and Cluster 4 IFN/STAT genes were positively enriched in pro-inflammatory M1-like macrophage genes induced by TNF and anti-correlated with the IL-4 driven M2 anti-inflammatory phenotype, indicating these two macrophage subsets are activated by pro-inflammatory factors (Fig. 1E, red v yellow bars). Conversely, Cluster 2 genes were negatively enriched for TNF-induced inflammatory response suggesting an anti-inflammatory phenotype, while the lack of enrichment with the IL-4 induced M2 state potentially indicates a novel tissue macrophage phenotype. The limited number of Cluster 3 positive markers did not significantly associate with any of the stimulated macrophage states.

As the most abundant cell type in the RA synovium (*28*) (Fig. S2A), synovial fibroblasts can evoke large shifts in macrophage gene expression profiles (*29*). Importantly, the combination of synovial fibroblasts together with TNF (TNF+Fib) generated a macrophage phenotype that aligned with Cluster 1 HBEGF^+^ macrophages more than the other stimuli (Fig. 1E). This was unlikely due to an additive effect that exacerbates the TNF response, as synovial fibroblasts and TNF can independently induce robustly opposing effects, for example in Cluster 4 genes (Fig. 1E). Thus, we posited that the abundant Cluster 1 HBEGF^+^ macrophages identified by single-cell analyses from affected human tissue are driven by the combination of tissue-specific factors from the resident synovial fibroblasts and inflammatory signals such as TNF found in the RA synovium.

Consistent with enrichment in RA, the CD14^+^ Cluster 1 HBEGF^+^ single-cells expressed high levels of classic inflammatory genes like *IL1B* and low levels of M2-associating genes such as *MERTK* (Fig. 1B, F)(*13*). However, Cluster 1 HBEGF^+^ cells were also distinctively high in the expression of genes such as *HBEGF* (Heparin Binding EGF Like Growth Factor) and *PLAUR* (plasminogen activator, urokinase receptor) (Fig. 1F), which have largely been reported as marking anti-inflammatory phenotypes or cells derived from a combination of pro- and anti-inflammatory triggers (*30, 31*). Considering this distinct program (Fig. 1G), we designate Cluster 1 macrophages ‘HBEGF^+^ inflammatory macrophages’.

Importantly, in an independent study using a droplet-based single-cell RNA-seq platform (Drop-seq) applied to five RA patient synovial tissues (Fig. S2A)(*28*), we have detected an abundant macrophage subset with considerable overlap of marker genes with the HBEGF^+^ inflammatory macrophages (Fig. S2B, 62%), including *HBEGF*, *PLAUR, IL1B* and *CREM* (Fig. S2C). Overlap with the other CD14^+^ single-cell clusters was less clear, potentially due to differences in the patient populations in the two studies. Nonetheless, through independent single-cell platforms and patient cohorts, synovial macrophages of the HBEGF^+^ inflammatory macrophage phenotype can be robustly detected in human tissue affected by RA.

### Tissue-resident synovial fibroblasts shape HBEGF+ inflammatory macrophages

To further define the HBEGF^+^ inflammatory phenotype, we compared the transcriptome of macrophages exposed to TNF and synovial fibroblasts versus TNF alone. In a transwell co-culture system where both cell types were exposed to TNF (24h, n=4 donors), synovial fibroblasts altered the expression of 3,709 macrophage genes (FDR adjusted p value < 0.1)(Fig. 2A). The majority of fibroblast-mediated effects opposed the direction of change brought on by TNF alone (Fig. 2A, 69% of fibroblast-effects anti-correlated with TNF effects, upper left + lower right). These changes included downregulation of TNF-inducible pro-inflammatory mediators such as *CFB*, *CXCL13*, *CCL8*, *MT1H*, *MMP2* and *SLAMF7* (Fig. 2A, upper left, genes not labeled). Alternatively, fibroblasts upregulated M2-like anti-inflammatory factors, otherwise suppressed by TNF, including *MRC1*, *MSR1*, and *TREM1* (Fig. 2A, lower right, genes not labeled). Pathway analysis indicated fibroblasts likely altered the metabolic state of TNF-treated macrophages, indicated by a collective suppression of factors involved in Oxidative Phosphorylation (Fig. S3A).

**Fig. 2.**
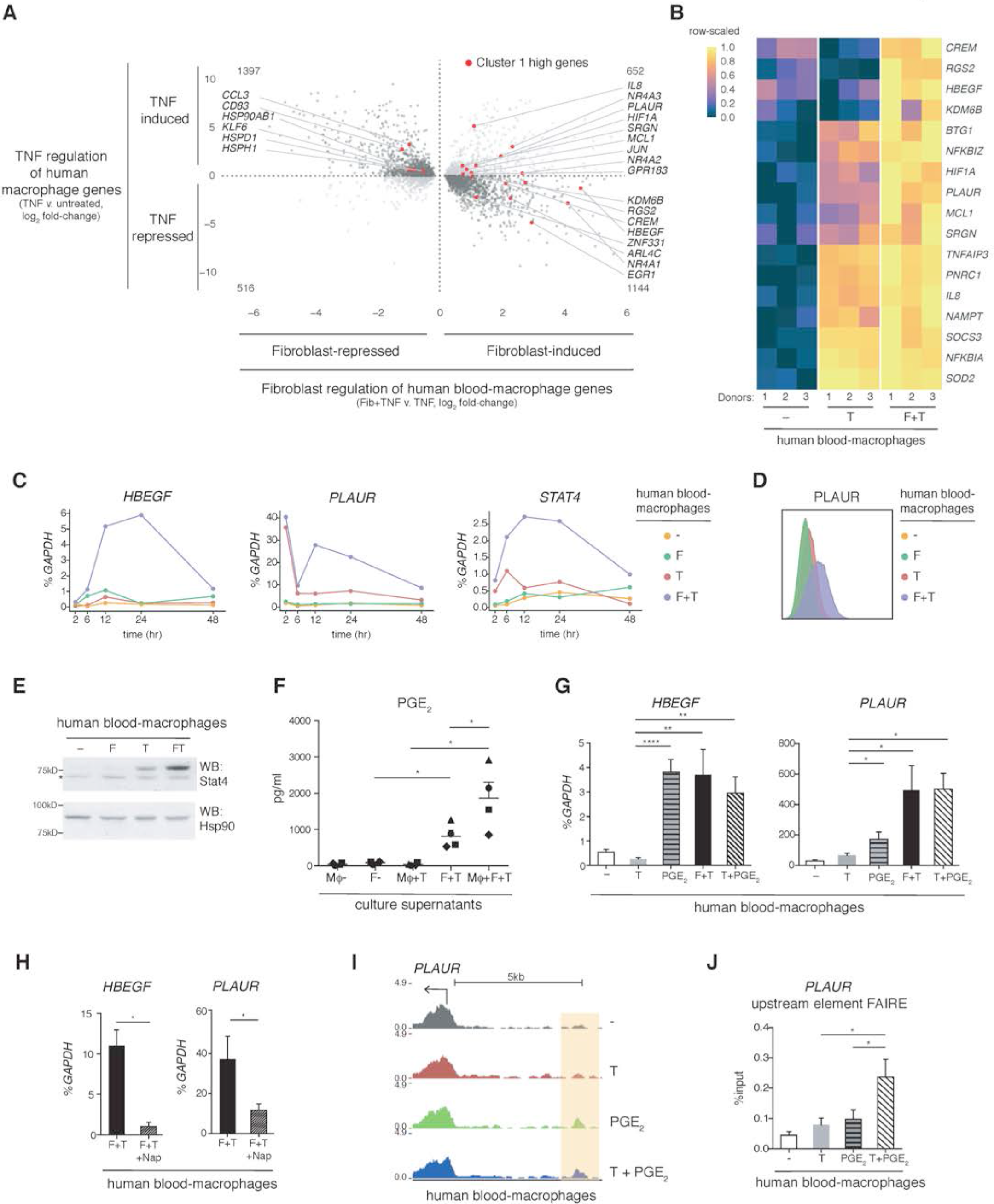
HBEGF^+^ inflammatory macrophages polarization by tissue fibroblasts. (**A**) Human blood-derived macrophage genes regulated by synovial fibroblasts and TNF (3,709 genes, FDR adjusted p < 0.1; n=4 donors). Expression changes plotted as log_2_-fold. x-axis plots Fibroblasts+TNF v. TNF; y-axis plots TNF v. untreated. HBEGF^+^ Cluster 1 single-cell markers labeled in red (expressed in >55% of cells, Bonferroni corrected p <10^−6^). (**B**) Expression of select synovial CD14^+^ Cluster 1 genes in the blood-derived macrophages exposed to TNF (T) or fibroblasts + TNF (F+T). n=3 donors. (**C**) qPCR of blood-derived macrophages overtime, plotted as percent (%) of GAPDH. Mean, standard error of the mean (SEM). Representative, n=4 donors. (**D**) *PLAUR* (CD87) surface protein detected by flow cytometry in blood-derived macrophages, 24h. Representative, n=3 donors. (**E**) Western blot of STAT4 in blood-derived macrophages, 24h. Representative, n=4 donors. *, non-specific band. Hsp90, loading control. (**F**) Prostaglandin E2 (PGE_2_) ELISA on supernatants from blood-derived macrophages (Mφ), 24h. n=4 donors. Mean with SEM. (**G**) qPCR of blood-derived macrophages, 24h. prostaglandin E2, (PGE_2_). n=8 donors. (**H**) qPCR of blood-derived macrophages, 24h. Nap, COX inhibitor naproxen (150nM). (**I**) ATAC-seq tracks from *PLAUR* gene promoter regions in blood-derived macrophages, 3h. Change in open chromatin, orange bar. (**J**) FAIRE-qPCR of open chromatin in region highlighted in subpanel **I**, % total input reported as mean SEM. n=4 donors. *, **, **** represent p<0.05, 0.01, 0.0001, respectively.

Despite the largely opposing effects of synovial fibroblasts on the macrophage TNF response, fibroblasts enhanced or maintained induction of a substantial portion of TNF-induced HBEGF^+^ inflammatory Cluster 1 genes, including *PLAUR* and *IL8* (Fig. 2A, upper right, red dots). Synovial fibroblasts also induced a portion of HBEGF^+^ inflammatory genes that were suppressed by TNF treatment alone, such as *HBEGF*, *RGS2* and *CREM* (Fig. 2A, lower right, red dots, and 2B). This indicates within synovial tissue, fibroblasts can also modulate macrophage polarization by inducing a portion of Cluster 1 genes that would otherwise be downregulated upon TNF exposure. The induction of *HBEGF* by synovial fibroblasts peaked around 12 hour and lasted over the course of days (Fig. 2C; qPCR representative of n=6 donors). The expression of *PLAUR* and the RA-associated transcription factor *STAT4*, while elevated transiently by TNF alone, was hyperinduced by synovial fibroblasts, resulting in a second elongated wave over days (Fig. 2C). Enhanced gene expression correlated with an increase at the protein level for cell surface PLAUR and intracellular STAT4 (Fig. 2D, E, respectively).

Cross-referencing the fibroblast-induced HBEGF^+^ macrophage profile with a panel of previously reported macrophage polarization states (*12*) demonstrated a strong correlation with macrophages treated with TNF and prostaglandin E2 (PGE_2_) and/or the Toll-like receptor 2 (TLR2) ligand Pam3-Cys (referred to as ‘TPP’ (*12, 32*)(Fig. S3B, red text). This strong correlation differed from the weaker or negative correlations with canonical M1 and M2 polarization states (induced by TNF or IFN-γ and IL-4 or IL-13, respectively)(Fig. S3B). From these data, we hypothesized that PGE_2_ mediated a considerable portion of the synovial fibroblast effect on macrophages. Indeed, TNF-stimulated synovial fibroblasts (Fig. 2F, F+T) produced large amounts of PGE_2_, whereas macrophages produced no detectable PGE_2_ irrespective of TNF exposure (Fig. 2F, MΦ and MΦ+T). Furthermore, induction of HBEGF^+^ genes such as *PLAUR* and *HBEGF* in blood-derived macrophages was recapitulated by exogenous PGE_2_ and TNF, with the fibroblast-mediated induction blocked by the COX enzyme inhibitor naproxen (Fig. 2G, H, respectively). The transcriptional induction of *PLAUR* expression associated with chromatin opening in the *PLAUR* promoter region, whereby TNF and prostaglandin synergized to open a new region, detected by ATAC-seq and verified by FAIRE-qPCR (Fig. 2I, J, respectively).

These data suggest that in the RA synovium, chronic exposure to the pro-inflammatory environment results in heightened synovial fibroblast production of prostaglandins that, together with inflammatory factors, drive macrophages towards a state distinct from classical M1 and M2 polarization. Unique transcriptional regulators, cell surface markers, metabolic pathways and chromatin modifications mark this HBEGF^+^ inflammatory state.

### HBEGF^+^ inflammatory macrophages promote fibroblast invasiveness

To examine how HBEGF^+^ inflammatory macrophages may impact RA pathology, we focused on their intercellular relationship with synovial fibroblasts. In an RNA-seq analysis, exposure to HBEGF^+^ inflammatory macrophages altered 855 synovial fibroblast genes (FDR adjusted p-value <0.1), including induction of *IL11*, *LIF*, *CSF3* (GCSF), *IL33*, *IL6* and *PLAU* (Fig. 3A). Pathway analyses revealed an upregulated EGFR response in the fibroblasts, observed with two gene sets (one induced by EGFR ligands and a second blocked by an EGFR inhibitor)(Fig. 3B). Similar to the gene expression changes, we also detected increased GCSF and IL-33 protein secretion in the presence of HBEGF^+^ macrophages (Fig. 3C). The increased molecular production of neutrophil chemoattractants and growth factors, such as GCSF, led us to investigate potential cellular effects of HBEGF^+^ inflammatory macrophages. We report neutrophil accumulation—a well-documented feature of RA joint fluid (*33, 34*) following co-culture of HBEGF^+^ macrophages with synovial fibroblasts (Fig. S4A). Lastly, we found the expression of GCSF and IL-33 was sensitive to EGFR inhibition (AG-1478 (AG))(Fig. 3D), further implicating EGFR involvement in this inflammatory macrophage-fibroblast crosstalk system and identifying an additional therapeutic avenue to target HBEGF^+^ macrophage driven inflammation.

**Fig. 3.**
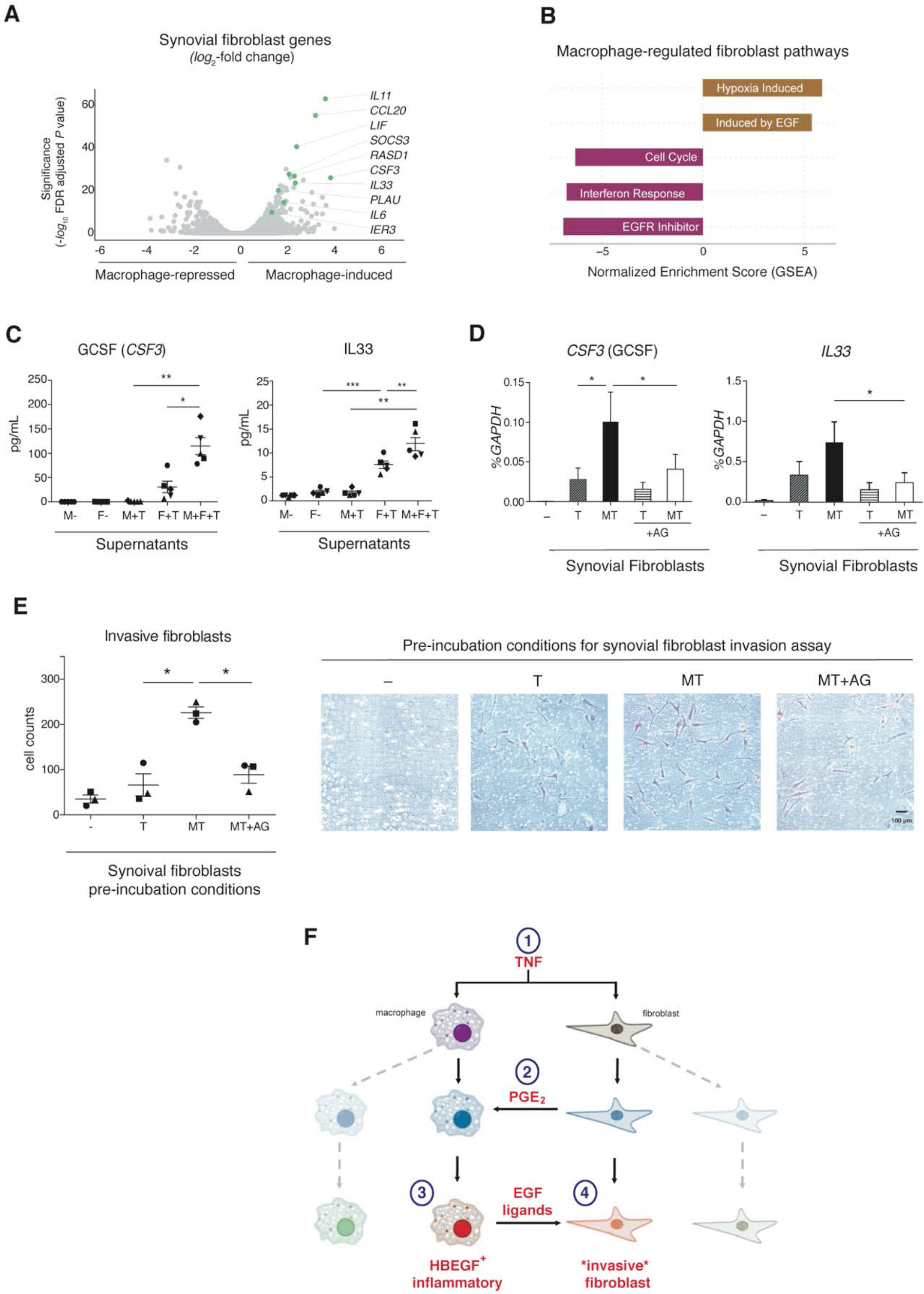
HBEGF^+^ inflammatory macrophages promote EGFR-dependent synovial fibroblast pathologic activity. (**A**) Human synovial fibroblast genes altered by blood-derived macrophages during a TNF response (885 genes differentially expressed), RNA-seq. 48h. n=2 donors. x-axis: *log*_2_ fold-change by macrophages. y-axis: significance as the −*log*_10_ FDR adjusted p-value. (**B**) Fibroblast pathways regulated by macrophages under TNF conditions (GSEA and Ingenuity Pathways Analysis (IPA). Macrophage-induced, brown. Macrophage-downregulated, maroon. (**C**) ELISA assay on supernatants of synovial fibroblast (F) cultures with or without macrophages (M) and TNF (T) for 48h. n=4 donors for both, reported as mean with SEM. (**D**) qPCR on fibroblasts, 32h. EGF receptor inhibitor, AG 1478 (AG). Mean, SEM. n=6 donors. (**E**) Synovial fibroblast Matrigel invasion assays, 18h after a 24h pre-incubation in with macrophages (M), TNF and the EGFR inhibitor (AG, 4 μM). n=3 donors. *, **, *** represent p <0.05, 0.01, 0.001 by paired Student’s t-test, respectively. (**F**) Unique inflammatory response arises for nearby synovial macrophages and fibroblasts, wherein [1] TNF induces [2] prostaglandin production by fibroblasts that together with TNF in macrophages drives [3] HBEGF+ inflammatory phenotype and EGF ligand production that then feeds back to induce [4] fibroblast invasive behaviors.

To further understand the EGFR-mediated gene expression in the synovial fibroblasts, we investigated the levels of EGF ligands in this system. Synovial fibroblasts expressed only trace amounts of seven EGF ligands with no change in response to TNF and HBEGF^+^ macrophages (Fig. S4B). In notable contrast, macrophages induced *HBEGF* and a second EGF ligand *EREG* (epiregulin) upon combined exposure of TNF and synovial fibroblasts (Fig. S4B). While *EREG* expression in the blood-derived macrophage system was higher than *HBEGF* (Fig. S4B), we have placed more emphasis on *HBEGF* as the RA patient synovial macrophages exhibit considerably higher expression of *HBEGF* (Fig. 1B). For EGF receptor expression, synovial fibroblasts expressed two subunits (EGFR and ERBB2), while neither blood-derived macrophages nor RA synovial macrophages expressed EGFR subunits (Fig. S4C, D). Considering these data, we posit synovial fibroblast EGFR responses correspond to production of EGF ligands by fibroblast-entrained inflammatory HBEGF^+^ macrophages.

EGFR signaling is a robust regulator of fibroblast motility (*35*) and in RA, synovial fibroblasts invade and destroy cartilage and bone (*36–39*). Thus, we next tested whether the heightened EGF response influenced fibroblast migration through extracellular matrix. Indeed, pre-incubation of fibroblasts with TNF and macrophages induced a pronounced increase in fibroblast invasiveness that was mitigated by EGF receptor inhibition (Fig. 3E, F). These data demonstrate that the HBEGF^+^ macrophage-dependent EGFR response may promote pathologic fibroblast-mediated and pannus-associated tissue destructive behaviors in RA joints.

### RA medications target HBEGF+ inflammatory macrophage polarization

We next examined how clinically effective RA medications impact the polarization of HBEGF^+^ inflammatory macrophages, using human blood-derived macrophages exposed to TNF and synovial fibroblasts. In comparison to macrophages exposed only to TNF, the presence of synovial fibroblasts substantially altered the impact of most medications (Fig. S5A). For example, the majority of auranofin targets were lost when macrophages were polarized towards the HBEGF^+^ phenotype (1,300 genes, 67%) (Fig. S5A, blue versus yellow). In contrast, several medications gained more than 1,000 gene targets in HBEGF^+^ inflammatory macrophages, including naproxen (Nap) and leflunomide (active metabolite A77) (Fig. S5A). Thus, the pronounced impact synovial fibroblasts confer onto macrophages includes alteration of drug responsiveness.

Several medications reversed a large portion of fibroblast-induced effects (~1,000 genes), suggesting the efficacy of these treatments could involve suppressed generation of HBEGF^+^ inflammatory macrophage in RA synovium (Fig. 4A, black). This included leflunomide (A77), dexamethasone (steroid), naproxen and Triple therapy (Tri, hydroxychloroquine + sulfasalazine + methotrexate)(Fig. 4A, black). As naproxen inhibits COX enzyme-mediated prostaglandin production and affected the majority (~80%) of genome-wide fibroblast-mediated effects (Fig. 4A and Fig. S5A-C), this further supports prostaglandins as a dominant fibroblast product driving HBEGF^+^ inflammatory macrophages. The drug-induced inhibition of HBEGF^+^ inflammatory macrophages, however, resulted in at least two functionally distinct states. Naproxen, for example, blocked the prostaglandin arm driving HBEGF^+^ inflammatory macrophages, but still permitted TNF polarization of the macrophages towards an M1-like pro-inflammatory phenotype, which presumably also functions pathologically in RA synovium (Fig. S5A-C). Medications like dexamethasone were capable of blocking large portions of both programs (Fig. S5A). This result highlights the importance of understanding the resulting phenotypes of macrophages targeted by medications, particularly in complex tissue settings where multiple factors could redirect macrophages towards a differing yet pathologic state.

**Fig. 4.**
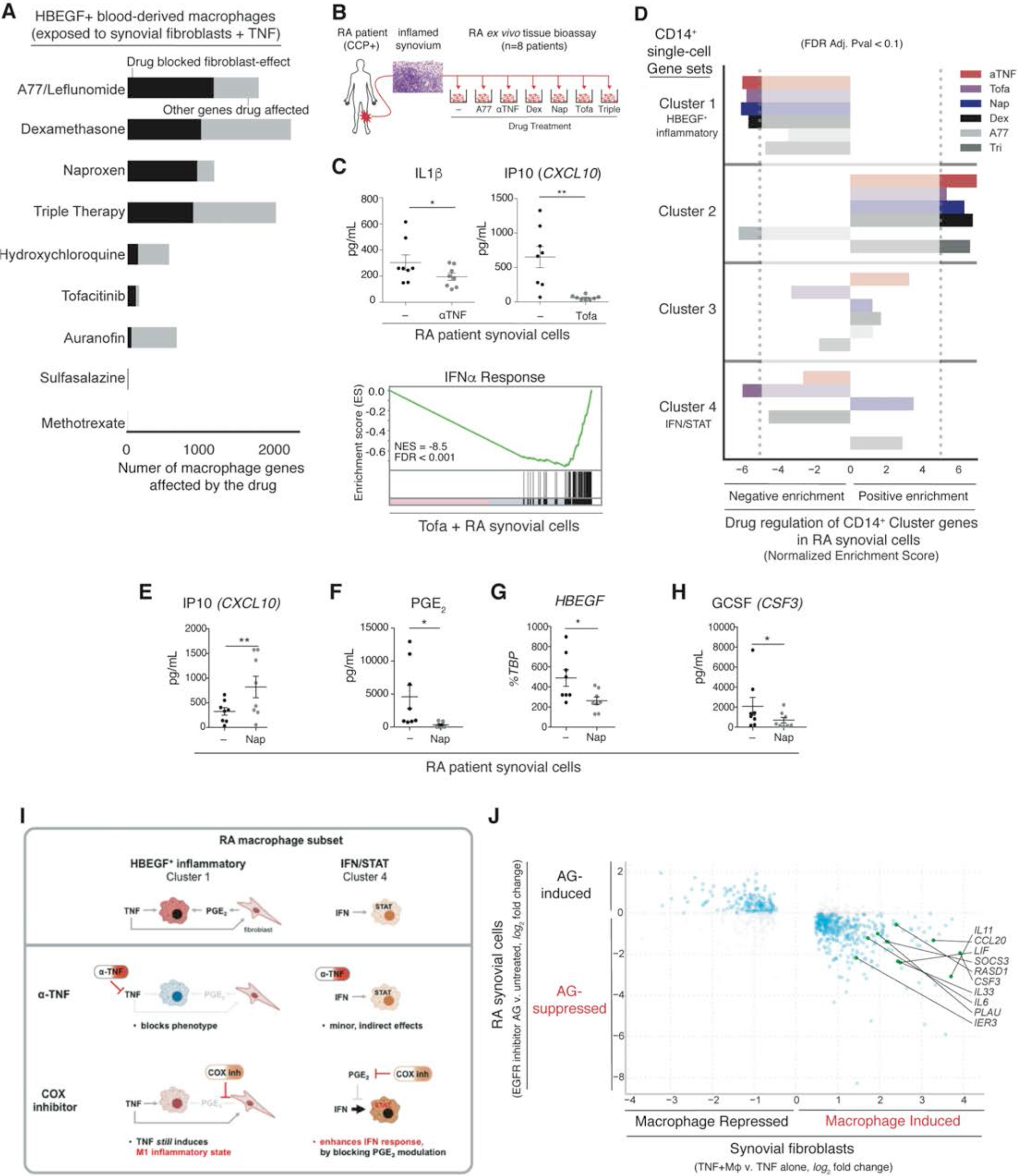
Clinically effective RA medications and an FDA-approved EGFR inhibitor target the HBEGF^+^ inflammatory macrophage-fibroblast crosstalk signatures in RA tissue. (**A**) Number of blood-derived macrophage genes affected by RA medications in the presence of TNF and synovial fibroblasts, 24h. Black, genes opposed by drug. Grey, all other genes regulated by drug. FDR adjusted p<0.1. n=2 to 4 donors. (**B**) RA patient synovial tissue *ex vivo* drug response assay using highly inflamed synovium. Dissociated cells were exposed in culture to a panel of medications, 24h. (**C**) ELISA on supernatants from RA tissue *ex vivo* assay. n=8 donors. *, **: p<0.05, 0.01, respectively by Wilcoxon signed-rank test. Lower panel: IFNα response upon tofacitinib exposure, bulk RNA-seq and GSEA. n=2 donors. (**D**) Normalized enrichment scores (GSEA) for drug-induced gene expression in the RA patient *ex vivo* assay for CD14^+^ single-cell Cluster markers. (**E**,**H**) ELISAs as described in c. (**F**) PGE_2_ ELISA on supernatants. Mean, SEM. n=7 donors. *, p<0.05, paired Student’s t-test. (**G**) qPCR on *ex vivo* synovial cells treated with naproxen, plotted as percent (%) of TBP. n=7 donors. Mean, SEM. *, p<0.05, paired Student’s t-test. (**I**) Schematic of differential effects of anti-TNFs and the COX inhibitor naproxen on the two RA-enriched synovial macrophage subsets. (**J**) RA patient *ex vivo* synovial expression changes with EGFR inhibitor AG-1478 (y-axis) compared to the synovial fibroblast gene regulation by macrophage and TNF (x-axis, data from Fig. 3A); n=2 donors. FDR adjusted p< 0.1, plotted as a *log*_2_ fold-change. Highlighted genes from Fig. 3A.

### RA medications target the HBEGF^+^ inflammatory macrophage-synovial fibroblast axis in patient samples

To examine the impact of therapeutically targeting HBEGF^+^ macrophages in RA joints, we directly assayed synovial tissue from RA patients with definitive RA diagnoses (*40, 41*) and extensive joint inflammation confirmed by histologic scoring (*42, 43*) (Table S1, n=8). Specifically, in an *ex vivo* tissue assay, synovium was dissociated into cell suspensions (*44*) and cultured with FDA-approved RA medications (Fig. 4B). Similar to the seminal assays that laid the precedent for anti-TNF therapies in RA (*45*), as well as the documented clinical responses (*46, 47*), exposure to anti-TNF antibodies (αTNF, adalimumab) suppressed production of the inflammatory mediator IL-1β (Fig. 4C). Furthermore in this assay, the JAK inhibitor tofacitinib imparted striking reductions in genome-wide IFNα responses and protein levels for the JAK/STAT target *CXCL10* (IP10) (Fig. 4C).

Notably in this tissue explant assay, anti-inflammatory therapies such as anti-TNF, tofacitinib and dexamethasone inhibited the HBEGF^+^ inflammatory synovial macrophage Cluster 1 gene set (Fig. 4D). Conversely, the panel of anti-inflammatory medications upregulated the Cluster 2 gene set (with one exception, A77/leflunomide) (Fig. 4D), further suggesting Cluster 2 cells possesses anti-inflammatory features, which also aligns with high levels of anti-inflammatory genes *MERTK*, *TGFBI* and *MARCO* (*48–50*)(Fig. 1F, Fig. S5D). Tofacitinib was the most potent inhibitor of the Cluster 4 ‘IFN/STAT’ gene set, further implicating a robust JAK/STAT response in this RA-associated macrophage subset (Fig. 4C). Interestingly, the JAK/STAT response in Cluster 4 was induced by naproxen, as shown by enrichment in Cluster 4 genes and an elevated *CXCL10*/IP10 protein production in naproxen-treated RA synovial cells (Fig. 4D-F). This correlated with reduced prostaglandins levels with naproxen and is in accord with the established inhibitory effects of prostaglandins on IFN responses (*51–53*).

To test for evidence of the crosstalk module we have identified between HBEGF^+^ inflammatory macrophages and synovial fibroblast EGFR responses (Figs. 2, 3), we further examined the effects of naproxen in our tissue explant system. In addition to blocking prostaglandin levels in the RA synovial explants and reducing *HBEGF* expression (Fig. 4G), naproxen also reduced GCSF (*CSF3*) secretion, indicating a decoupling of the macrophage-fibroblast circuit (Fig. 4H). Importantly, as mentioned above and in Fig. S5 and B, naproxen does not interfere with TNF-mediated effects and thus despite dampening prostaglandin levels in the tissue assay (Fig. 4F), naproxen treatment did not suppress IL-1b protein production in the *ex vivo* synovial cell assay (data not shown). TNF blockade, however, was effective in blocking *HBEGF* expression, as well as, IL-1b and GCSF protein levels (Fig. 4C, D) and thereby derails generation of both the HBEGF^+^ inflammatory macrophages-synovial fibroblast axis and pro-inflammatory M1-like macrophages (Fig. 4I).

To directly assay EGFR-dependent fibroblast gene expression changes consistent with HBEGF^+^ inflammatory macrophage influence, we introduced an EGFR inhibitor into the *ex vivo* assay. Importantly, the EGFR inhibitor reversed the majority of the synovial fibroblast gene expression effects elicited by HBEGF^+^ inflammatory macrophages (77%), including a suppression of *CCL20, CSF3, LIF, IL33, IL11*, and *PLAU* (Fig. 4J, 3A). Together, these data provide evidence within patient tissues for the crosstalk module we identified between HBEGF^+^ inflammatory macrophages and tissue-resident fibroblasts and furthermore indicates blockade of EGFR responses may provide a non-immunosuppressive therapeutic approach for RA.

### Discussion

Challenges in treating autoimmune and inflammatory disorders have led to strategies including administering medications sequentially until one is found to improve clinical symptoms. In an effort to improve *a priori* knowledge of which medication will be most effective, we have taken into consideration the complexity of cellular interactions in affected tissues and identified a population of HBEGF^+^ inflammatory macrophages in inflamed RA joints and delineated their deterministic tissue-resident and disease-specific factors. Furthermore, we have identified medications that successfully interfere with the generation of these macrophages and the support they provide towards fibroblast invasiveness, which contributes to irreversible joint destruction in RA.

The crosstalk mechanisms and functional output between macrophages and fibroblasts in the RA tissue environment differ from other pathologic states driven by these two cell types. In fibrosis, macrophages and fibroblasts drive excessive collagen deposition blocking tissue function in part by static physical barriers rather than invasion and erosion (*2, 54, 55*). We find in RA, fibroblast products such as prostaglandins combine with chronic inflammatory signals such as TNF to polarize macrophages into a state that is distinct from inflammatory M1, anti-inflammatory wound healing M2 and pro-fibrotic TGFβ-expressing macrophages. The RA-enriched HBEGF^+^ inflammatory macrophages produce a defined subset of inflammatory products such IL-1 and the EGF growth factors HBEGF and epiregulin. As HBEGF^+^ inflammatory macrophages subsequently induce invasiveness in synovial fibroblasts (Fig. 3F), we classify them as ‘pro-invasive macrophages’.

While leukocyte-derived growth factors have emerged as critical components in tissue homeostasis and repair (*1, 56*), our data uncover a pathologic correlate. In the RA tissue environment, macrophage-produced EGF ligands lead to EGFR-dependent synovial fibroblast invasiveness and GCSF production concomitant with enhanced neutrophil accumulation (Fig. 3)—dominant features of RA joint pathology (*33, 34, 37*). In inflammatory arthritis *in vivo* models, disease progression is driven by EGF ligands, EGF receptor activity and the enzyme iRhom2, which mediates release of both TNF and HBEGF from immune cells (*57–60*). Our data provide relevance to these findings in human disease and a mechanistic understanding of the cellular crosstalk program involving growth factors in RA joints. Furthermore, considerable evidence implicates TNF and HBEGF as pathologic drivers of kidney disease in the autoimmune condition lupus (*61, 62*). Thus, linked TNF and EGFR responses may be a unifying and targetable feature in tissues affected by disparate autoimmune conditions.

Using perturbations relevant to human disease, namely FDA-approved medications, we have detailed the disruption of intercellular interactions in patient samples and the resulting phenotypes, thereby gaining insights into the complex consequences of medication in RA joint tissue. In particular, our data has yielded insights into a longstanding question of why anti-inflammatory COX inhibitors known as NSAIDs are not “disease-modifying” in RA (*15*). Our data suggest NSAIDs block the prostaglandin-mediated arm in HBEGF^+^ inflammatory polarization, but still permit macrophage TNF responses (Fig. 4I, Fig. S5B-C). Thus, NSAID therapy in RA likely redirects HBEGF^+^ inflammatory macrophages towards a classic pro-inflammatory M1-like phenotype, which would presumably perpetuate inflammation albeit through a different pathway. In that regard, NSAIDS may best be used in combination with medications like anti-TNFs, in order to target two arms of HBEGF^+^ inflammatory macrophage polarization. However COX inhibition by naproxen also proved problematic in the other RA-enriched macrophage phenotype, inducing the Cluster 4 ‘IFN/STAT’ response (Fig. 4I), consistent with robust suppression of IFN responses by prostaglandins (*51, 53*). Thus for two RA-enriched macrophage phenotypes, NSAIDS are permissive of pathologic responses. These data speak to a greater need in understanding tissue microenvironment factors that polarize macrophages and how in the presence of therapeutics the intercellular communication networks are rewired and subsequently repolarize macrophages into states that either resolve or perpetuate pathology.

Along with the seminal synovial explant assay that incited the use of TNF therapies in RA (*45*), our work impels the implementation of *ex vivo* assays to better understand and treat patients with autoimmune and inflammatory disorders. Specifically, in addition to detecting genome-wide established targets of anti-TNF and tofacitinib therapies, our RA patient *ex vivo* synovial tissue bioassay provides a human- and disease-relevant system that unmasked the interconnectivity and drug responsiveness of the synovial macrophage-fibroblast interaction we identified. Human tissue-based therapeutic testing could offer guidance in relevant therapeutic choices based directly on the response of the unique cellular composition of a patient’s tissue. For RA, this could be accomplished with the expanding use of synovial biopsies (*21, 22, 63*) and ultimately over time with the identification of circulating biomarkers that correlate with tissue-based assays. Lastly, effective blockade of the macrophage-induced fibroblast response in the RA tissue *ex vivo* assay with an EGFR inhibitor developed for cancer warrants testing of this as a new treatment direction, particularly as it could represent a non-immunosuppressive treatment option.

## MATERIAL and METHODS

### Study Design

The experimental objectives for this study involved directly analyzing primary human tissues to understand how tissue and disease factors influence macrophage responses in the autoimmune disease RA. This study included the use of tissue samples from patients treated for RA and OA, in addition to healthy human blood donors. Inclusion and exclusion criteria for the arthritis patients were based on standard clinical diagnostic criteria. Perceived outliers in sequencing datasets remained in the analyses upon application of surrogate variable analysis by svaseq 3.26.0. Human donor blood-derived macrophage biologic replicates used for various assays depended on the robustness of the response for each type of assay and therein the amount of variability seen across donors, whereby for robust assay responses typically ~n=4 donors were used but for assays with higher variability up to n=8 donors were used.

### Patient recruitment and CD14+ synovial cell sorting for RNA-seq

The multicenter RA/SLE Network of the Accelerating Medicines Partnership (AMP) consortium enrolled individuals meeting the ACR 2010 RA classification according to protocols approved by the institutional review board at each site (*22, 44*). Synovial tissues were collected from ultrasound-guided biopsies or joint replacement surgery and viably frozen in Cryostor CS10 cryopreservation media (Sigma-Aldrich). At a central processing site, tissues were dissociated and cells were FACS sorted (BD FACSAria Fusion) into fibroblast, macrophage, B cell and T cell populations. Macrophages were sorted based on CD14+CD45+ cell surface expression. For bulk CD14+ synovial cell population RNA-seq, ~1,000 cells were sorted directly into RLT buffer (Qiagen). For CD14+ synovial single-cell RNA-seq, ~100 live cells per patient were individually plated and lysed in 384 well plates.

### Bulk RNA-seq of sorted CD14+ synovial cell population

Full-length cDNA and sequencing libraries were generated using Illumina Smart-Seq2 protocol (*64*). Libraries were sequenced on a MiSeq System (Illumina) to generate 35 length base pair, paired-end reads.

### Single-cell RNA-seq of sorted CD14+ synovial cells

Single-cell RNA-seq was performed on sorted macrophage and synovial fibroblast using the CEL-Seq2 protocol (*65*) in 384 well plates. Libraries were sequenced on a HiSeq 2500 (Illumina) in Rapid Run Mode to generate 76 length base pair, paired-end reads.

### CD14+ single-cell RNA-seq read alignment and differential gene expression analysis

RNA-seq reads were aligned with STAR version 2.5.2b (*66*) to the hg19 reference genome. Transcript levels were quantified as Counts Per Million using the GENCODE Release 24 annotation. The single cell gene expression matrix was clustered based on a canonical correlation analysis (CCA) methodology (*22*). Briefly, highly variable genes identified from single-cell RNA-seq and bulk RNA-seq datasets were integrated based on genes that maximized the correlation between the two datasets. The correlated canonical variates were then used to construct a nearest neighbor network thereby generating clusters that are verified to be present in the bulk data. Using the four clusters identified from the CCA method, the top ten canonical coordinates were used to generate a Euclidean distance matrix for tSNE visualization using a perplexity parameter of 40. Positive and negative cluster markers were identified using the Wilcoxon rank sum test with a Bonferroni correction for multiple testing.

### CD14+ bulk RNA-seq read alignment and differential gene expression analysis

Reads were aligned and quantified with STAR version 2.4.2a (*66*) against the GRCh38 genome and GENCODE Release 27 annotation, respectively. Differential expression analyses and batch-correction were performed using DESeq2 (*67*) version 1.18.1 and svaseq (*68*) version 3.26.0, respectively.

### Pathway analysis

Gene lists were processed for GSEA(*69*) version 3.0 by taking the inverse of the FDR adjusted p-value for each gene and multiplying it by the sign of the log2 fold change relative to the baseline conditions. GSEA was run under the “pre-rank” mode with 1,000 permutations for each of the gene sets available in MSigDB and ImmunSigDB Version 5.2. Additional pathway analysis was performed using QIAGEN’s Ingenuity® Pathway Analysis (IPA®, QIAGEN Redwood City, www.qiagen.com/ingenuity). Reported pathways were referenced as follows with the MSigDB systematic name in parentheses: TNF-a Signaling via NFkB (M5890), Interferon-g Response (M5913), Interferon-a Response (M5911), Inflammation (M5932), MYC Targets (M5926, M5928), Oxidative Phosphorylation (M5936), Translation (M11989), Cell Cycle (M543), Induced by EGF (M2613), Hypoxia Induced (M5891), Interferon Responsive Genes (M9221), EGFR Inhibitor (M16010), and KEGG Ribosome Pathway (M189). Gene sets derived from the CD14+ synovial single-cell RNA-seq markers were composed of up to 500 genes that exhibited >0.5 log2 fold change (positive marker) or <−0.5 log2 fold change (negative marker genes) relative to all other clusters, sorted by their fold change.

### Independent RA arthroplasty cohort analyzed by droplet-based single-cell RNA-seq

In an independent analysis, we collected synovial tissue from five RA patients consented under HSS RA Studies (IRB#2014-317 and 2014-233) during arthroplasty and synovectomy procedures. Tissues were dissociated into single cell preparations and all cells were run through Drop-seq protocol and sequenced on a HiSeq 2500 (Illumina)(*28*). Following cell and gene filtering(*28*), we applied Seurat version 2.3.0 to generate PCA based single cell clusters, which were labeled based on cell-type markers. 20,031 single cells were visualized using the t-SNE implementation in Seurat using a perplexity parameter of 20 and 13 principal components. After identifying a macrophage cluster consistent with our previous results (4,212 single cells), we re-applied Seurat and identified distinct subpopulations and visualized in t-SNE space.

### Cell culture for human blood-derived macrophages and synovial fibroblasts

Human CD14+ monocytes were purified from leukocyte preparations purchased from the New York Blood Center and differentiated into blood-derived macrophages for 1–2 d in 10 ng/ml M-CSF (PeproTech) and RPMI 1640 medium (Life Technologies)/10% defined FBS (Hyclone). Cells were stimulated with 20 ng/ml recombinant human TNF (PeproTech). Drug treatments were administered 15 minutes after TNF exposure to both the top and bottom wells of the transwell system to achieve the stated final concentrations. After suspending in DMSO according to the company’s instructions, auranofin (Sigma Aldrich) was added to cells at 500 nM, A77 1726 (Santa Cruz) was added at 50 mM, Tyrphostin AG 1478 (Sigma Aldrich) at 4 μM, GW 627368X (Cayman Chemical) at 10 μM, sulfasalazine (Sigma Aldrich) at 3 μM, hydroxychloroquine sulfate (Sigma Aldrich) at 50 μM, methotrexate (Cayman Chemical) at 110 μM, tofacitinib citrate (Cayman Chemical) at 1μM_ENREF_61, and Pam3CSK4 (InvivoGen) at 1000 μg/mL. Dexamethasone (Sigma Aldrich), and prostaglandin E2 (Sigma) were first suspended in absolute ethanol and then added to cells at 100 nM, and 280 nM, respectively. Naproxen was suspended in RPMI supplemented with 10% FBS and then added to cells at a final concentration of 100 uM. Anti-TNF was provided as adalimumab and was added to cells at a final concentration of 50 μg/mL.

Human synovial fibroblasts derived from de-identified synovial tissues of RA patients undergoing arthroplasty (HSS IRB#14-033)_ENREF_60. Dissociated cells were plated in aMEM based media up to 10 days, washing the media numerous times to remove dying blood cell components. Synovial fibroblasts at passages 4-6 were used for experiments. The diagnoses of RA were based on the ACR 2010 criteria. For Transwell culture experiments, synovial fibroblasts adhered to polyester chambers with 0.4-μm pores (Corning) and were suspended above the wells containing macrophages, with a ratio of fibroblasts to macrophages of 1:16 based on the size of the cells and their coverage of the culture well surface. The number of donors used for each experiment is listed in the figure legend and refers to unique donors for both the blood-derived macrophages and for the synovial fibroblast lines.

### RNA-seq for human blood-derived macrophages and synovial fibroblasts

Total RNA was first extracted using RNeasy mini kit (Qiagen). Tru-seq (non-stranded and PolyA selected) sample preparation kits (Illumina) were then used to purify poly-A transcripts and generate libraries with multiplexed barcode adaptors. Single-end libraries were multiplexed, pooled, and sequenced using the Single Read Clustering with 100 cycles for 3 lanes and 50 cycles for 6 lanes on an Illumina HiSeq 4000. Paired end libraries were multiplexed, pooled and sequenced using the Paired End Clustering protocol with 51×2 cycles sequencing for 4 lanes on an Illumina HiSeq 2500. Sequencing was performed by the Weill Cornell Medical College Genomics Resources Core Facility. RNA-seq read alignment, quantification, differential testing, and pathway analysis was performed as previously described.

### Real-time PCR

RNA obtained using RNAeasy Mini kit (Qiagen) with DNAse treatment was reverse transcribed into cDNA (Fermentas) and analyzed by real-time quantitative PCR (Fast SYBR Green;Applied Biosystems) on a 7500 Fast Real-Time PCR System (Applied Biosystems). Gene-specific primer sequences were as previously described_ENREF_63 or listed in the Table below. Expression levels were normalized to GAPDH or TBP.

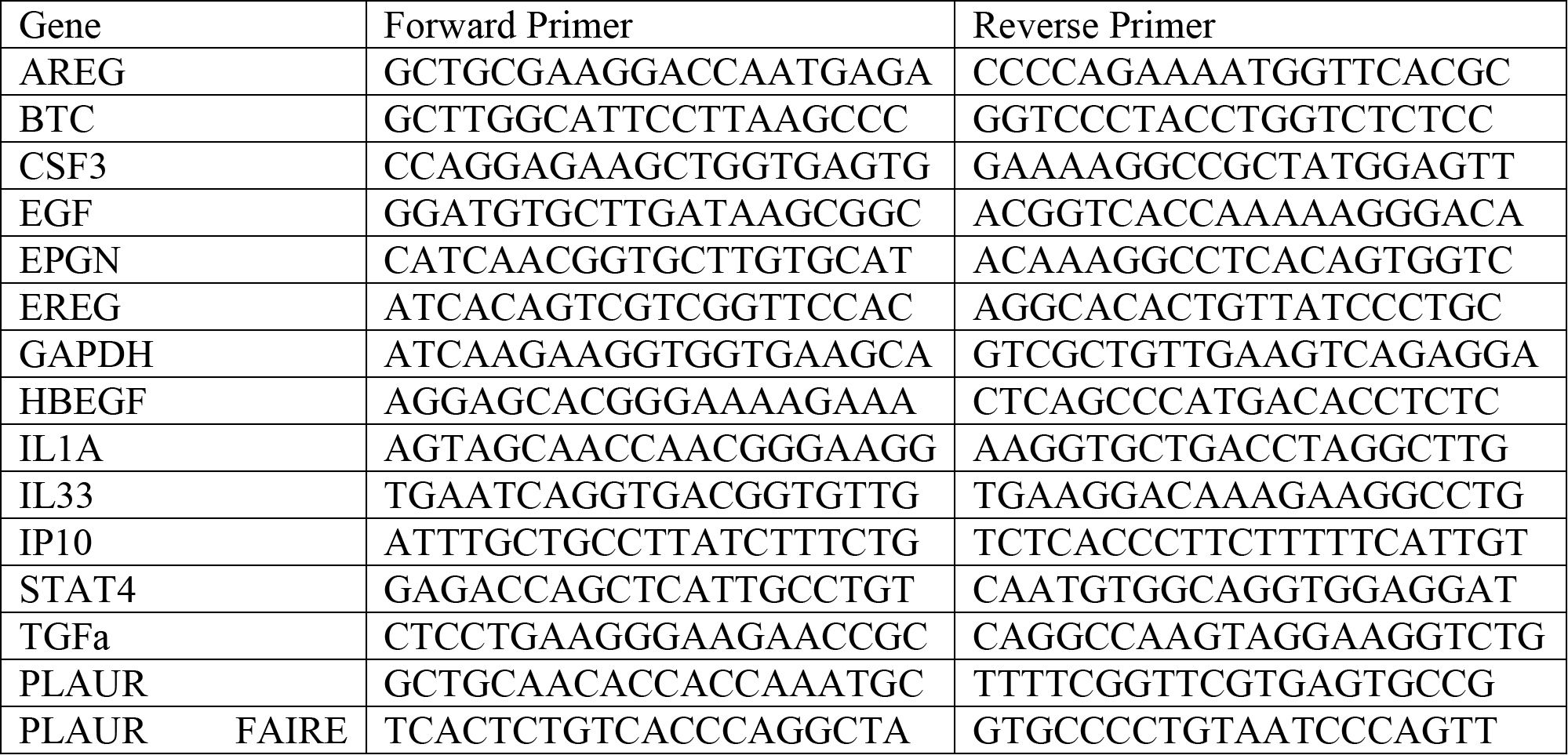

### Western blots on human blood-derived macrophages

Western blot analyses using a STAT4 (Santa Cruz) and HSP90 (Cell Signaling Technology) antibody were performed using standard procedures with the additional step of adding Pefabloc (Sigma-Aldrich) to macrophage cultures before cell lysis to prevent STAT protein degradation.

### Flow cytometry on human blood-derived macrophages

Blood-derived macrophages were resuspended in 100uL FACS buffer (PBS with 1% FBS) and stained with FITC-conjugated CD87/PLAUR (VIM5-Miltenyi Biotec), acquired by FACSCanto (BD Biosciences) and analyzed using FlowJo (Tree Star, Inc.) software.

### Assay for Transposase-Accessible Chromatin (ATAC)-Seq

ATAC-seq was performed as previously described(*70*). Human blood-derived macrophages were treated for 3 hours with 20ng/ml TNF (Peprotech) and/or 280nM PGE2 (Sigma-Aldrich). 50,000 cells were washed in PBS, lysed (lysis buffer: 10 mM Tris-HCl, pH 7.4, 10 mM NaCl, 3 mM MgCl2 and 0.1% IGEPAL CA-630) and centrifuged immediately at 500g for 10 min to obtain nuclei. The pellet was resuspended in the transposase reaction mix (25 μl 2× TD buffer, 2.5 μl transposase (Illumina) and 22.5 μl nuclease-free water) and incubated at 37°C for 30 min. DNA was purified using a Qiagen MinElute kit and library fragments amplified for 13 cycles using 1X NEB next PCR master mix and custom Nextera PCR primers(*70*). The libraries were purified using Agencourt AMPure XP PCR Purification kit (Beckman Coulter) and single-end sequenced on a HiSeq 2500 (Illumina).

### FAIRE qPCR

FAIRE experiments were performed as previously described (*71*). Chromatin was crosslinked by treating cells with 1% formaldehyde for 7 min and the reaction quenched with 0.125 M glycine for 5 min. Cells were washed with cold PBS and scraped, followed by a second wash. Fixed cells were lysed in buffer LB1 (50 mM HEPES-KOH, pH 7.5, 140 mM NaCl, 1 mM EDTA, 10% glycerol, 0.5% NP-40, 0.25% Triton X-100, and protease inhibitors) for 10 min. Pelleted nuclei were resuspended in buffer LB2 (10 mM Tris–HCl, pH 8.0, 200 mM NaCl, 1 mM EDTA, 0.5 mM EGTA, and protease inhibitors) and incubated on a rotator for 10 min. The nuclei were pelleted and lysed in buffer LB3 (10 mM Tris–HCl, pH 8.0, 100 mM NaCl, 1 mM EDTA, 0.5 mM EGTA, 0.1% Na-deoxycholate, 0.5% N-lauroylsarcosine, and protease inhibitors). Chromatin was sheared using a Bioruptor Pico device (Diagenode). A total of 10% of sonicated nuclear lysates were saved as input. Phenol-chloroform-purified nuclear lysates and de-crosslinked input DNA were used for qPCR analysis using specific primers for the upstream region. Chromatin accessibility is displayed relative to total input.

### Prostaglandin E2 EIA Kit

Culture supernatants were collected from the culture after 24 hours, diluted 1:50 in plain RPMI, and prostaglandin concentrations measured using the Prostaglandin E2 EIA Kit-Monoclonal (Cayman Chemical). ELISA plates were read on Varioskan Flash Multimode Reader (ThermoFisher Scientific).

### Neutrophil viability assay

Supernatants were collected from cultures of human macrophages and synovial fibroblasts after 24 hours and frozen at −80 °C. Whole blood from healthy human subjects was collected into heparin coated tubes. Neutrophils were isolated with the EasySep Direct Human Neutrophil Isolation Kit (Stemcell Technologies) with “The Big Easy” EasySep Magnet. In a 12-well plate, 600,000 neutrophils were plated in 800 μL of supernatant for 24 hours. Neutrophil viability was measured using Muse Annexin V and Dead Cell Assay Kit on a Muse Cell Analyzer mini-flow cytometer (EMD Millipore).

### Fibroblast invasion assay

Human synovial fibroblasts were plated in transwells with macrophages as in the transwell experiments. AG 1478 (Sigma Aldrich) was added at 4 μM for 32 hours. Fibroblasts were trypsinized and re-plated in 500μL plain alpha-MEM with 10 ng/ml M-CSF at 0.1 × 106 cells per well into 24-well Corning BioCoat Matrigel Invasion Chambers. Macrophages were resuspended in 750 μL plain alpha-MEM and seeded underneath invasion transwells in the appropriate conditions. AG 1478 was added at a concentration of 4 μM; after 18 hours, the fibroblasts were fixed for 10 minutes in ice-cold methanol and stained using crystal violet. Invasive fibroblast numbers were quantified via light microscopy.

### Human RA synoviocyte cultures

For synoviocyte cultures, RA patient synovial tissue was obtained from patients consented into the HSS FLARE study (IRB# 2014-233). Tissues were digested with Liberase TL (100 μg/mL, Roche) and DNaseI (100 μg/mL, Roche) for 15 minutes and passed through three 70 μM cell strainers. Cells were then suspended in 1 mL RBC Lysis Buffer (gift of J. Lederer, BWH) for 3 minutes followed by addition of RPMI/10%FBS/1%Glutamine to quench the reaction. Disaggregated synoviocytes were plated in RPMI/10%FBS/1%Glutamine at 0.2 × 106 in 96-well plates. Cells were treated with drugs at aforementioned concentrations for 24 hours. Supernatants and RNA were collected for Luminex experiments and quantitative PCR, respectively. For RNA-seq, the samples were multiplexed in eight samples per lane, 50 cycles, single-end reads, with Truseq (Illumina) for library prep and a HiSeq 4000 (Illumina) ran in the Weill Cornell Medical College Genomics Resources Core Facility. RNA-seq read alignment, quantification, differential testing, and pathway analysis was performed as previously described(*29*).

### Luminex

Supernatants were collected from macrophage-fibroblast cultures or synoviocyte cultures at 48 hours. Customized Luminex panels were ordered from R&D Systems. Protein concentrations were read by a MAGPIX from EMD Millipore using xPONENT 4.2.

### Statistical analysis

Data are shown as mean and standard error unless stated otherwise. Two-tailed paired t-tests were performed for human sample derived qPCR and ELISA data using GraphPad Prism version 5.04 for Windows. Luminex multiplex ELISA experimental data was tested for normality by the Shapiro-Wilks test with a significance threshold of p < 0.05 and was found to not follow a Gaussian distribution. Subsequent statistical analysis was thus performed by Wilcoxon signed rank tests, a non-parametric method, using GraphPad Prism. Single-cell RNA-seq clusters were identified using a Canonical Correlation Analysis (CCA)(*22*). Markers for different clusters were determined by Bonferroni corrected Wilcoxon rank sum tests implemented in Seurat version 2.3.0. Visualization of intersecting sets was performed using UpSetR version 1.3.3(*72*). Testing for differentially expressed genes from bulk RNA-seq count data was performed using DESeq2 version 1.18.1 and surrogate variable analysis was performed using svaseq version 3.26.0, both run on the R version 3.3.2. All RNA-seq significance levels are reported as False Discovery Rate (FDR) adjusted p-values. Spearman and Pearson correlation analysis was performed using the seaborn statistical data visualization package version 0.8.1 run on Python version 3.6.4. Statistical significance of pathway analysis reported as the Normalized Enrichment Score and FDR adjusted p-values determined from testing of 1,000 permutations from GSEA version 3.0. Pathway z-scores for upstream regulatory analyses were performed using Ingenuity® Pathway Analysis (IPA®, QIAGEN). All data transformations for visualization purposes, replicate numbers, and statistical significance thresholds are reported in the Methods and Figure Legends.

### Data Availability

RNAseq data have been deposited to database of Phenotypes and Genotypes (dbGaP) with the accession numbers phs001340.v1.p1 and phs001529.v1.p1 as well as the Gene Expression Omnibus (GEO) database with the accession numbers GSE57723, GSE95588, and GSE100382. The CD14+ synovial single-cell data generated by the AMP consortium is housed at http://immport.org with the study accession numbers SDY998 and SDY999.

## Supplementary Materials

**Fig. S1.** Gene markers for synovial CD14^+^ single-cell clusters.

**Fig. S2.** Identification of synovial HBEGF^+^ inflammatory macrophages in an independent RA patient study.

**Fig. S3.** The transcriptome of human blood-derived macrophage exposed to synovial fibroblasts and TNF compared with published pathways and macrophage polarization phenotypes.

**Fig. S4.** Synovial fibroblasts express EGF receptors while HBEGF+ inflammatory macrophages express two EGF ligands.

**Fig. S5.** RA medications impose highly variable effects depending on inflammatory macrophages depending on the presence of synovial fibroblasts.

**Table S1.** Baseline Characteristics.

**Table S2.** Participants Characteristics.

## Acknowledgements

We thank the HSS orthopedic surgeons (particularly Dr. M. Figgie), rheumatologists, clinical research coordinators (particularly R. Cummings, M. McNamara and S. Mirza), and the consented HSS patients; Drs. P. Gulko and T. Laragione (Mt. Sinai) for technical guidance on synovial fibroblast invasion assays; R. Yuan, D. Oliver, and E. Giannopoulou for assistance with sequencing data; and C. Blobel and H. Hang for critically reading the manuscript. We also acknowledge R. Satija, W. Stephenson and A. Bulter of the New York Genome Center for their previous work on synovial tissue single-cell analysis using Drop-seq technology (*28*). We acknowledge the Accelerating Medicines Partnership (AMP): RA/SLE Network for the multi-site and large-scale collection of arthritis patient synovial tissue samples, cell sorting and RNA sequencing. We thank the AMP clinicians, scientists administrators and patients. AMP is a public–private partnership (AbbVie, Arthritis Foundation, Bristol-Myers Squibb, Lupus Foundation of America, Lupus Research Alliance, Merck Sharp & Dohme, NIH, Pfizer, Rheumatology Research Foundation, Sanofi, and Takeda Pharmaceuticals International) created to develop new ways of identifying and validating promising biologic targets for diagnostics and drug development.

## Author Contributions

DK analyzed sequencing data, prepared figures, and edited the manuscript; JD performed experiments and analyzed data; IC performed experiments and prepared figures; FZ analyzed sequencing data; KW designed experimental pipelines and performed experiments; DK designed experimental pipelines and performed experiments; US performed experiments; CR performed experiments; AMP recruited patients and performed experiments; EFD scored the histology slides; MBB designed and supervised experimental pipelines; VPB and SMG collected clinical data and samples; SR and GR supervised sequencing data analyses; LBI jointly conceived the project, acquired funding and edited the manuscript; LTD jointly conceived the project, acquired funding, wrote the manuscript, and supervised the research experiments and analyses.

## Data and materials availability

Accession codes and databases for sequencing data can be found in the Materials and Methods section.

