## Supplemental Materials for "HBEGF^+^ macrophages identified in rheumatoid arthritis promote joint tissue invasiveness and are reshaped differentially by medications"

### for

**Supplementary Materials**

**Fig. S1.** Gene markers for synovial CD14$^{+}$ single-cell clusters.

**Fig. S2.** Identification of synovial HBEGF$^{+}$ inflammatory macrophages in an independent RA patient study.

**Fig. S3.** The transcriptome of human blood-derived macrophage exposed to synovial fibroblasts and TNF compared with published pathways and macrophage polarization phenotypes.

**Fig. S4.** Synovial fibroblasts express EGF receptors while HBEGF+ inflammatory macrophages express two EGF ligands.

**Fig. S5.** RA medications impose highly variable effects depending on inflammatory macrophages depending on the presence of synovial fibroblasts.

**Table S1.** Baseline Characteristics.

**Table S2.** Participants Characteristics.

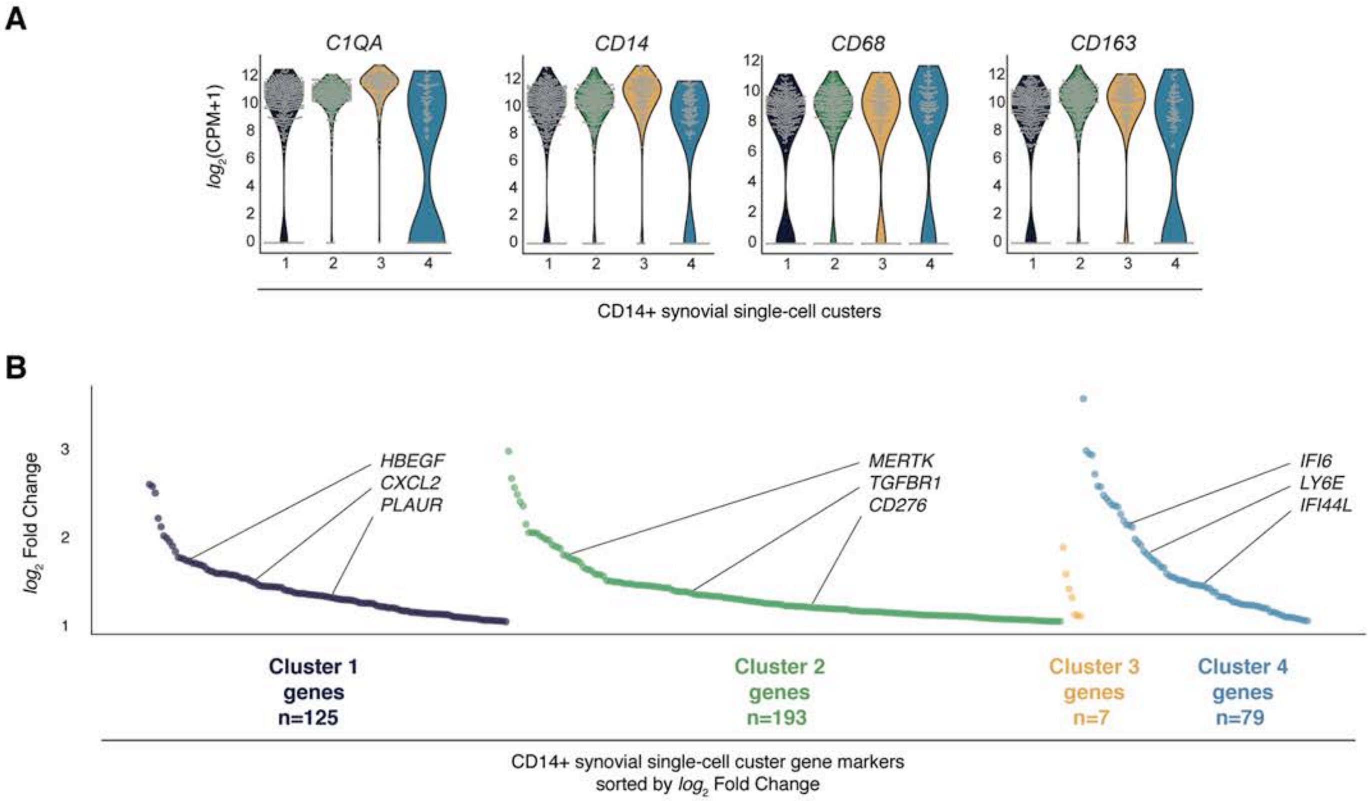


**Fig. S1. Gene markers for synovial CD14⁺ single-cell clusters.** (**A**) Violin plots for the expression of monocyte-macrophage marker genes in the four synovial CD14⁺ single-cell clusters from Fig.1. Expression represented as the $log_2$ counts per million + 1(CPM+1). (**B**) Scatter plots for synovial CD14⁺ single-cell cluster markers. Markers were selected based on a $log_2$ fold-change greater than 1 relative to all other clusters (y-axis) and are sorted in descending order for each cluster (x-axis).

Kuo et al.
Fig. S2

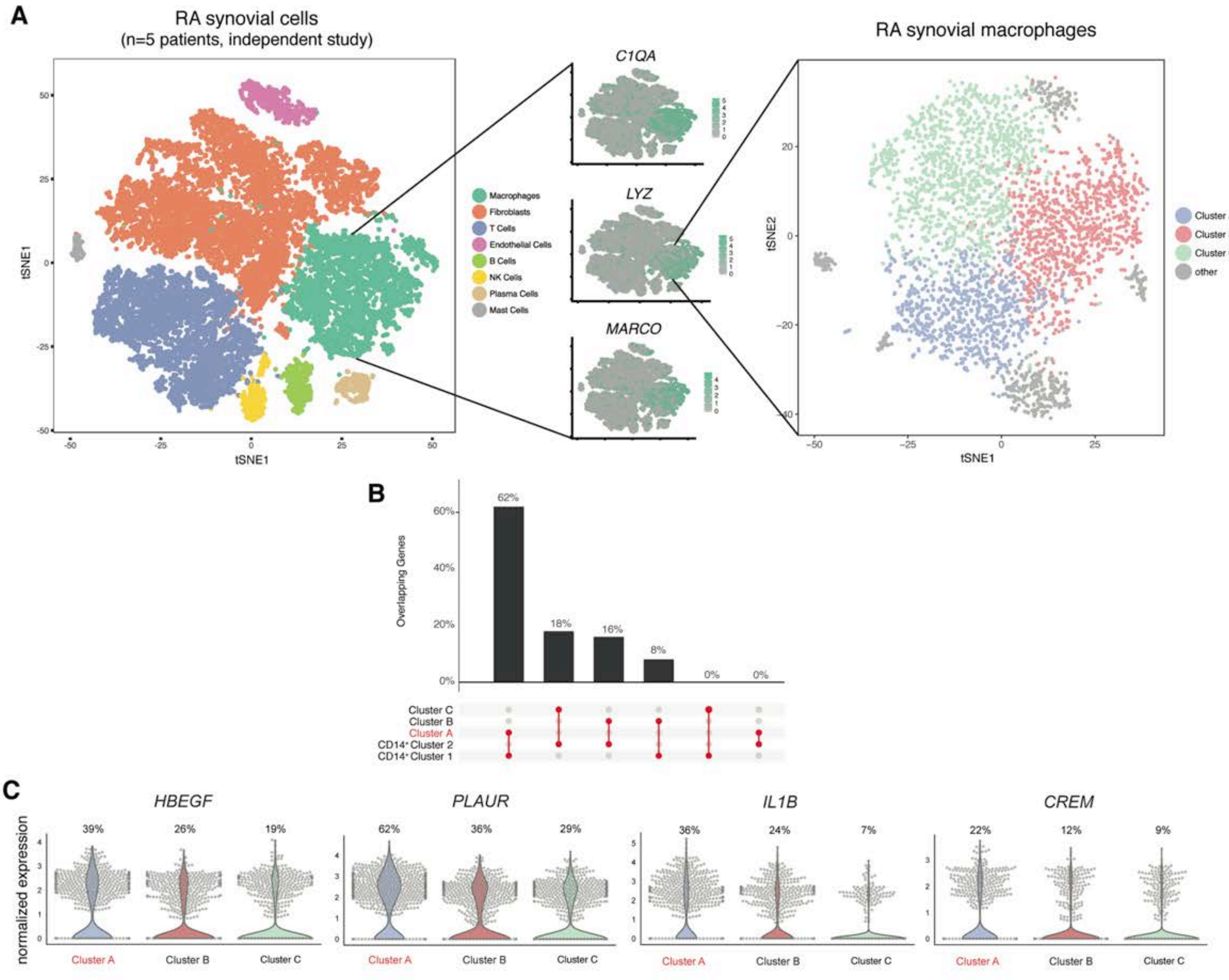


**Fig. S2. Identification of synovial HBEGF⁺ inflammatory macrophages in an independent RA patient study.** (**A**) From an independent RA patient synovial tissue study (Stephenson et al. Nat Comm 2018), tSNE plots of single-cell RNA Drop-seq data clustered by cell type (left, 20,031 cells) and macrophage specific cells re-clustered (right, 4,212 cells). Center panel depicts cells expressing three macrophage lineage markers in light green. n=5 RA. (**B**) Barplot depicting the percentage of overlapping markers between the CD14⁺ synovial cell clusters from Fig.1 (CD14+ Cluster 1-2) and macrophage clusters from Stephenson et al. visualized in a (Clusters A-C). The top markers from the CD14⁺ Clusters 1-2 (n=50) were compared to the top makers (n=300) from Clusters A-C. (**C**) Violin plots for Clusters A-C using marker genes for HBEGF+ inflammatory macrophages. Percentage values represent the percentage of cells for whichsome level of expression for that gene was dectected (i.e. non-zero cells).

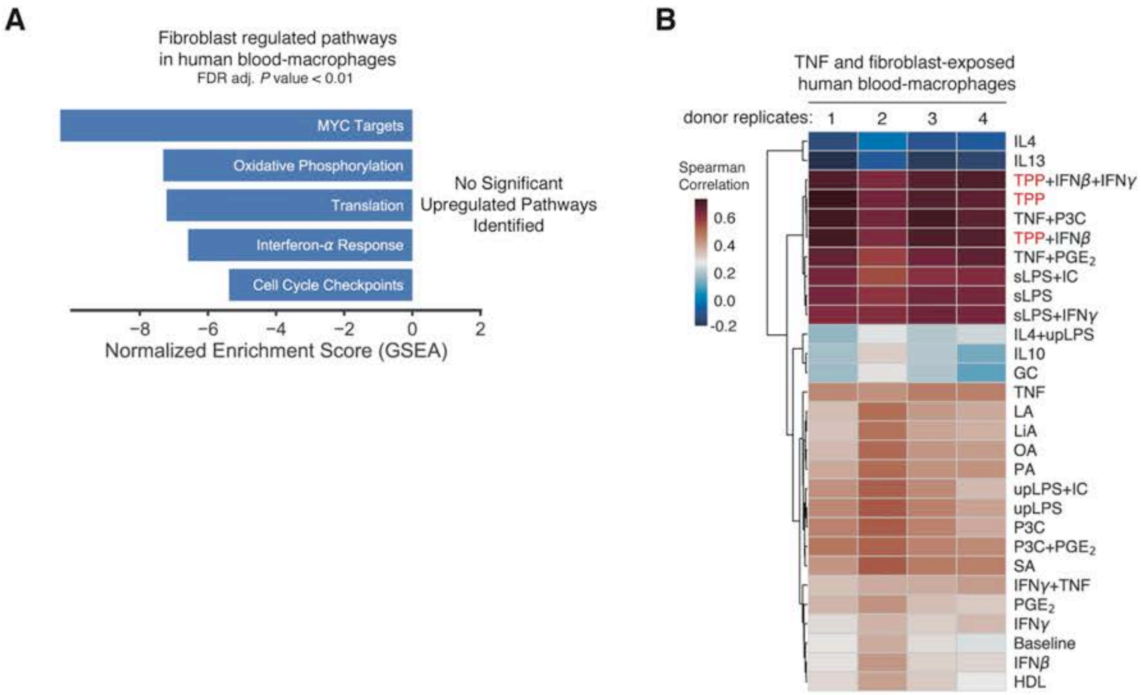


**Fig. S3. The transcriptome of human blood-derived macrophage exposed to synovial fibroblasts and TNF compared with published pathways and macrophage polarization phenotypes.** (**A**) GSEA and IPA (Ingenuity Pathway Analysis) results identified macrophage pathways downregulated by synovial fibroblasts in the presence of TNF (i.e. pathways with negative enrichment scores). No significantly upregulated pathways were identified by this analysis. (**B**) Heatmap of Spearman rank correlation values using the top 250 most variable and overlapping genes for macrophages stimulated under a variety of polarization conditions (rows; microarray expression data from Xue et al.) and four donor human blood-derived macrophages cultured with fibroblasts and TNF (columns; RNAseq normalized expression data).

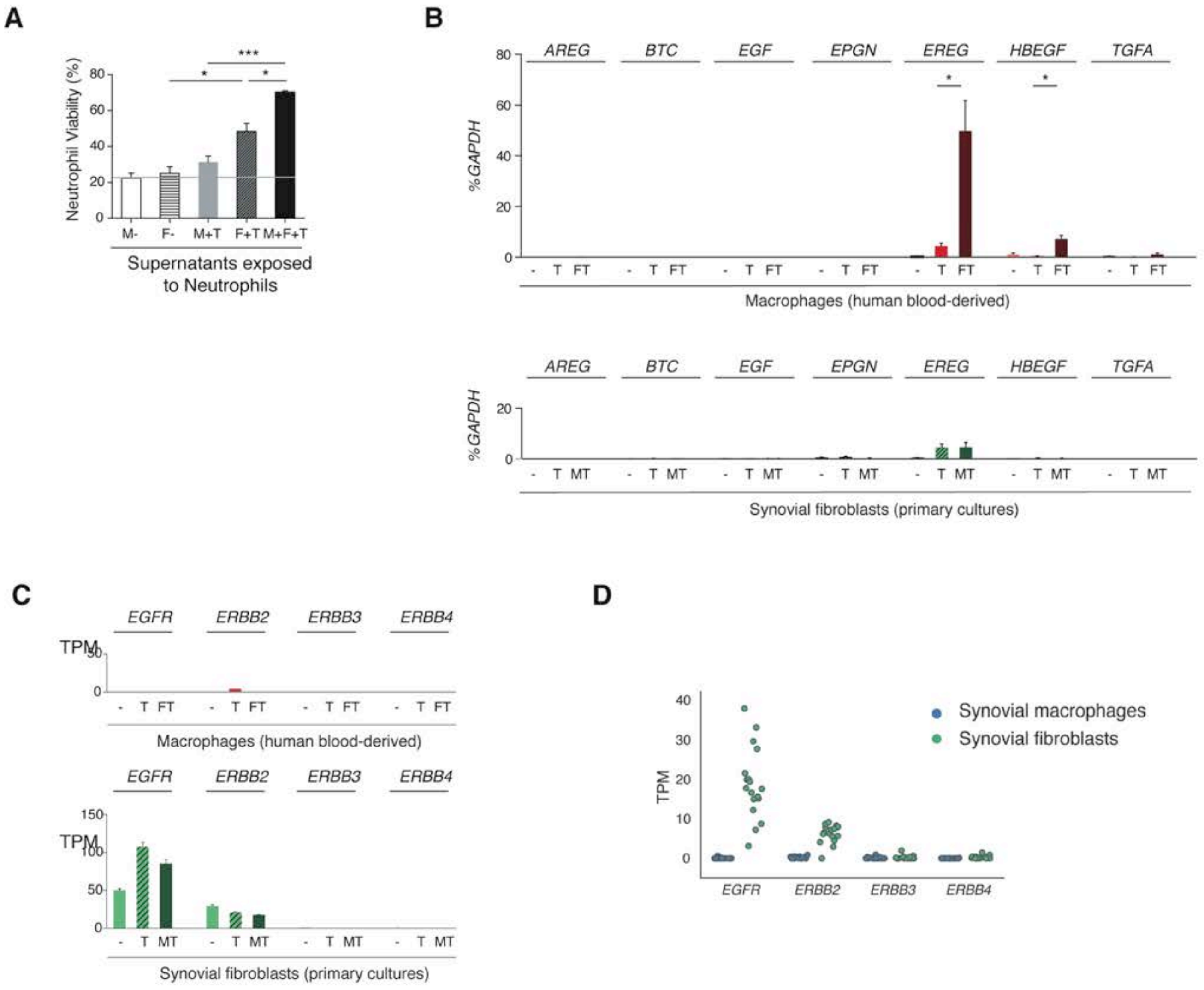


**Fig. S4. Synovial fibroblasts express EGF receptors while HBEGF+ inflammatory macrophages express two EGF ligands.** (A) Flow cytometry detection of neutrophil apoptotic and cell death markers were used to calculate viability after 24h incubation with supernatants from macrophage and fibroblast mono- and co-cultures. n=4 unique donors for all cell types reported as mean with standard error of the mean. (B) Quantitative PCR analyses of fibroblasts for seven EGF ligands cultured alone (—), in the presence of TNF (T), or with macrophages and TNF (MT) for 24h. Gene expression levels are plotted as percent (%) of GAPDH; error bars represent standard error, n=4 unique donors for each cell type. (C) Quantitative PCR analyses of fibroblasts for four subunits of the EGF receptor cultured alone (—), in the presence of TNF (T), or with macrophages and TNF (MT) for 24h. Gene expression levels are plotted as percent (%) of GAPDH; error bars represent standard error, n=4 unique donors for each cell type. *, **, *** represent p-value<0.05, 0.01, 0.001 by paired Student's t-test, respectively. (D) Stripplot of RA patient low-input RNAseq gene expression (measured as transcripts per million) of EGF receptors from synovial macrophages (blue dots, n=20) and synovial fibroblasts (green dots, n=18).

Kuo et al.
Fig. S5

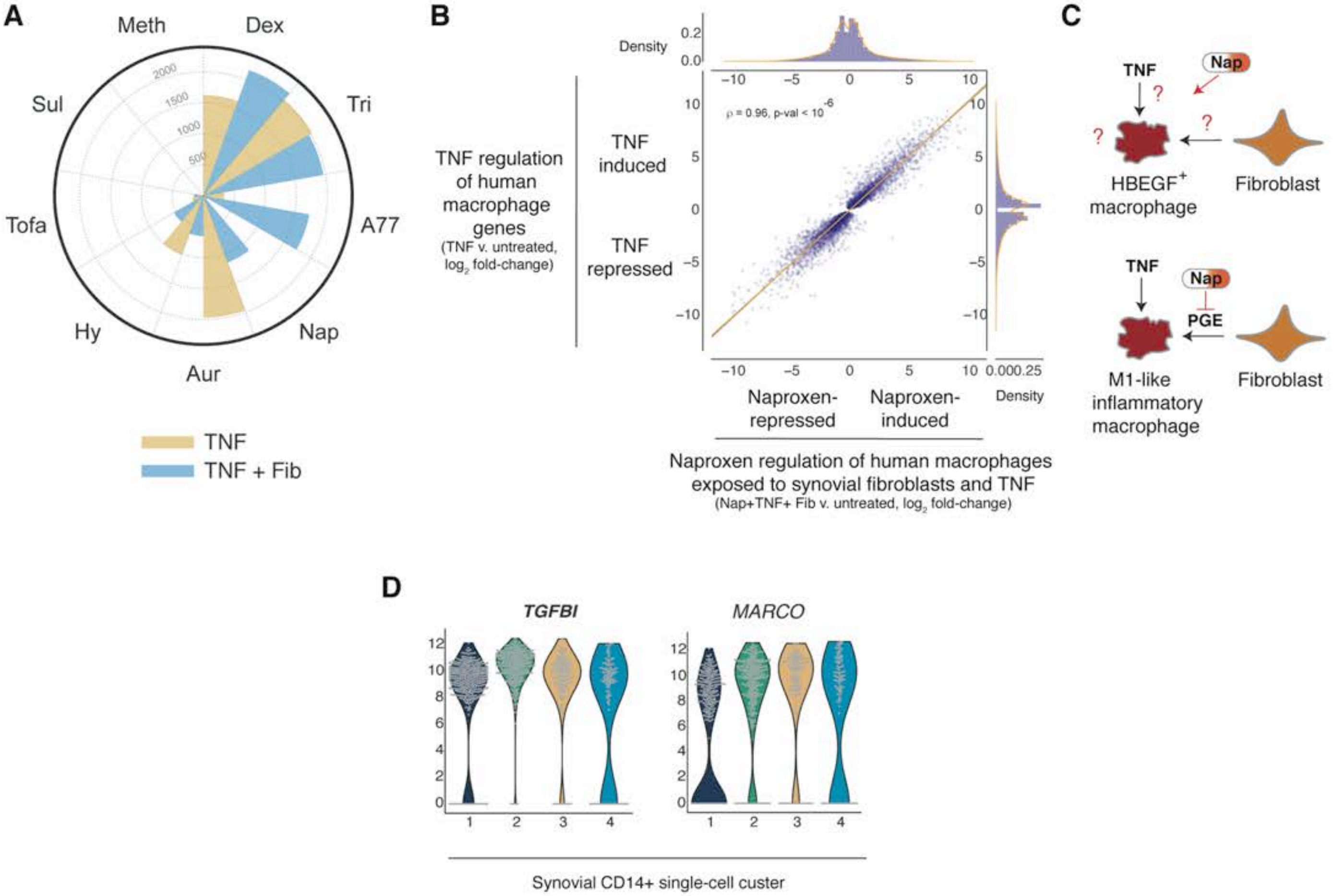


**Fig. S5. RA medications impose highly variable effects depending on inflammatory macrophages depending on the presence of synovial fibroblasts.** (**A**) Radar chart plotting the total number of drug regulated genes (FDR adjusted Pvalue < 0.1) from ten different drugs (spokes demarcated by black dotted lines). The yellow bar represents the total number of significant drug regulated genes in macrophages treated with or without drug after TNF stimulation (TNF). The blue bar represents the total number of significant drug regulated genes in macrophages treated with or without drug after TNF stimulation and fibroblast coculture (TNF+Fib). Drugs are ordered clockwise by the total number of TNF+ Fib drug regulated genes. (**B**) Scatter plot of TNF regulated genes relative to untreated human macrophages (n=6,613; FDR adjusted p-value < 0.1). The y-axis plots the $\log_2$ fold-change of the TNF v. untreated comparison andthe x-axis plots the $\log_2$ fold-change of naproxen regulation of macrophages stimulated with TNF and exposed to fibroblasts v. untreated. The orange line represents the regression between the $\log_2$ fold-change values (Person's $\rho = 0.96$, p-value $< 10^{-6}$). (**C**) Graphical representation of naproxen effect on $HBEGF^{+}$ inflammatory macrophages. Naproxen could impact TNF regulation, fibroblast-regulation and/or other responses in $HBEGF^{+}$ inflammatory macrophages (top panel, red question marks). Data in subpanels a and b suggest naproxen targets almost exclusively the fibroblast-mediated effects on $HBEGF^{+}$ inflammatory macrophages. As a Cox-inhibitor, this suggests naproxen blocks fibroblast prostaglandin production and thereby the polarization of $HBEGF^{+}$ inflammatory phenotype, but does permit TNF polarization towards an M1 inflammatory phenotype. (**D**) Violin plots of selected marker genes from CD14+ single-cell RNAseq represented by the *$\log_2$* counts per million (CPM) (y-axis) across the four identified clusters (x-axis).

**Table I. Baseline Characteristics ($N = 8$)**

| Characteristic | Average Value | Range or Percentage |
|---|---|---|
| Age (years) | 66.1 | 61 – 79 |
| Female | 5 | 62.5% |
| Race: White | 5 | 62.5% |
| BMI ($kg/m^2$) | 28.1 | 20.9 – 33.2 |
| Duration Dx (years) | 8.6 | 0.1 – 21.1 |
| Duration Sx (years) | 9.5 | 0.2 – 22.4 |
| ESR | 36 | 20 – 54 |

**Table II. Participant Characteristics**

| | RA Diagnosis | Serology | | Synovial Tissue Histology | | RA Medication | |
|---|---|---|---|---|---|---|---|
| MS ID | Criteria | RF Result Interpretation | anti-CCP Interpretation | Lymphocytic Infiltration | Lining Hyperplasia | DMARD | Biologic |
| Patient 1 | Meets BOTH | Positive | High Positive* | +++ | ++++ | Yes | Yes |
| Patient 2 | Meets BOTH | Positive | High Positive* | +++ | ++++ | Yes | No |
| Patient 3 | Meets BOTH | Positive | High Positive* | ++++ | +++ | No | No |
| Patient 4 | Meets BOTH | Positive | High Positive* | ++++ | +++ | Yes | Yes |
| Patient 5 | Meets BOTH | Positive | High Positive* | ++++ | ++++ | Yes | Yes |
| Patient 6 | Meets BOTH | Negative | Negative | ++++ | ++++ | Yes | Yes |
| Patient 7 | Meets 2010 | Negative | Negative | ++ | ++++ | No | Yes |
| Patient 8 | Meets 2010 | Negative | High Positive* | +++ | ++++ | Yes | No |

RA Diagnosis: BOTH indicates 2010 ACR/EULAR Criteria and 1987 ACR Criteria;
Serology: High Positive* signifies >3x ULN;
Histology: Symbols correspond to pathologist scored gradient from mild (+) to severe (++++);
